# Hidden Challenges in Evaluating Spillover Risk of Zoonotic Viruses using Machine Learning Models

**DOI:** 10.1101/2024.04.25.591033

**Authors:** Junna Kawasaki, Tadaki Suzuki, Michiaki Hamada

## Abstract

Machine learning models have been deployed to assess the zoonotic spillover risk of viruses by identifying their human infectivity potential. However, the scarcity of comprehensive datasets poses a major challenge, limiting the predictable range of viruses. Our study addressed this limitation through two key strategies: constructing expansive datasets across 26 viral families and developing new models leveraging large language models pre-trained on extensive nucleotide sequences. Our approaches substantially boosted our model performance. This enhancement was particularly notable in segmented RNA viruses, which are involved with severe zoonoses but have been overlooked due to limited data availability. Furthermore, models trained on data up to 2018 displayed strong generalization capability for viruses emerging post-2018. Nonetheless, we also found remaining challenges in alerting the zoonotic potential of specific viral lineages, including SARS-CoV-2. Our study elaborates on the models and datasets for predicting viral infectivity and highlights the unresolved issues to fully exploit machine learning in preparing for future zoonotic threats.

## Introduction

Because zoonotic viruses pose a significant threat to human health, monitoring animal viruses with the potential for human infection is crucial^1–3^. Despite advances in metagenomic and metatranscriptomic research revealing vast viral genetic diversity in animals^4,5^, the evaluation of viral phenotypes, such as infectivity, transmissibility, pathogenesis, and virulence, requires substantial human efforts due to the lack of high-throughput methods. Human infectivity, an important phenotypic characteristic relevant to zoonotic viral spillover and subsequent emerging diseases, remains largely unvalidated for most viruses. To address these challenges, machine learning models for predicting human infectivity using viral genetic features as inputs have been developed^2,3,6–10^. These models may help determine priority viruses for further virological characterization.

Given our limited understanding of the mechanisms driving viral infectivity, unsupervised feature extraction from viral genetic sequences is necessary. Large language models (LLMs), pre-trained on extensive genetic data, have achieved state-of-the-art performances in various genotype-to-phenotype tasks^11^. Pre-trained LLMs are expected to capture context-like rules in nucleotide sequences, allowing the construction of high-performance models even with limited labeled data. Furthermore, leveraging LLMs for viral infectivity prediction tasks can potentially uncover previously indiscernible patterns crucial for predicting viral infectivity. Thus, such unsupervised feature extraction could improve model performance and deepen our understanding of molecular mechanisms underlying viral infectivity.

While existing models have demonstrated high performance, several gaps remain in the model evaluation scheme^2,12,13^. First, the absence of standardized datasets limits the uniform comparison of model performances. Second, previous evaluations may have overestimated predictive performance because the dataset was occupied by viruses that did not match the purpose of predicting viral infectivity in humans, such as bacteriophages^6,7,10^. Third, given the successive emergence of novel zoonotic viruses, models should preferably be generalizable to unknown viruses (i.e., emerging after the model construction). Addressing these challenges in model development is essential for their real-world application in public health management.

Here, we aimed to improve viral infectivity prediction by (i) curating datasets covering various viral families and (ii) developing new models by leveraging LLMs. Our models outperformed existing models for most viral families. Nonetheless, we also found the general limitations in current machine learning models: the difficulty in alerting the human infectious risk in specific zoonotic viral lineages. This study provides a comprehensive benchmark for viral infectivity prediction models and highlights the remaining challenges of fully leveraging machine learning in preparedness for upcoming zoonotic threats.

## Results

### Constructing comprehensive large-scale datasets for 26 viral families

We developed models for each viral family by training with pairs of viral sequences and their host information (**Fig. 1A**). While the Virus-Host Database^14^, a curated source for virus-host relationships, has been mainly used for training dataset in previous studies^6,7,10^, its dataset composition does not perfectly align with the purpose of the models: predicting the zoonotic potential of viruses. For example, this database (version 2019/01/31, used in^7^) included 13,396 viral sequences, but only 77.6% were associated with eukaryotic hosts (**Supplementary Table 1**), which may lead to overestimating model performance owing to the presence of easy-to-predict viruses, such as bacteriophages.

**Figure 1:**
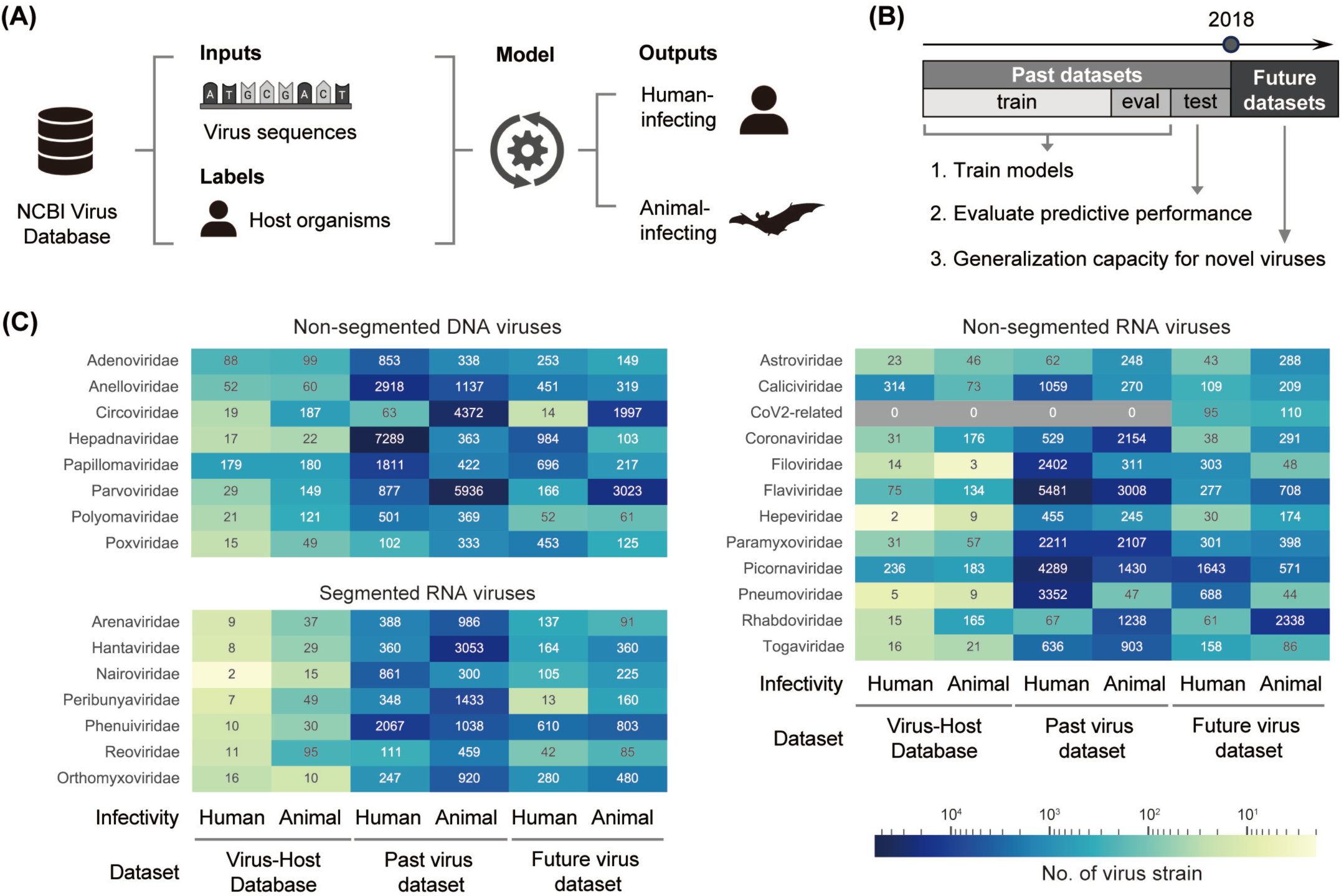
Training and evaluation of predictive models for viral infectivity. **a** Dataset preparation and model training (**see Methods**). **b** Data splitting to reflect read-world data availability. **c** Number of virus strains in our datasets and those in the Virus-Host Database^14^ mainly used in previous studies^6,7^.

To construct models more suitable for human-infecting viruses, we collected data from the NCBI Virus database^14^ for 26 viral families that include key human-infecting pathogens (**details in Methods, Supplementary Table 2-4**). Because the human infectivity of many viruses has not been directly validated, the infectivity label (i.e., human-infecting or animal-infecting) for each viral strain was defined based on the host information from which the viral sequences were collected (**see Discussion**). Our datasets included viruses associated with 1,476 vertebrate species and 535 arthropod species.

Eventually, our curated datasets offered a substantial increase in available data, approximately 29-times more than that of the Virus-Host Database^14^ (**Fig. 1C**). Notably, the Virus-Host Database contained >20 human-infecting viral strains for 15 of the 26 viral families, potentially overlooking important pathogens during model training and evaluation. By contrast, our comprehensive datasets encompassed at least 50 human-infecting viral strains for each viral family in the past virus datasets, establishing a valuable resource for developing predictive models for viral infectivity.

### Evaluation strategy and comparative models

To assess the capability of the predictive models for generalization to newly emerged viruses, we divided the viral data into two datasets: (i) a past virus dataset comprising sequences identified up to the end of 2017 for model training and (ii) a future virus dataset for evaluating the generalization capability toward viruses discovered post-2018 (**Fig. 1B**).

To address the potential shortage of labeled data for viral infectivity, we utilized LLMs pre-trained on extensive genetic sequences^11^. We fine-tuned two pre-trained Bidirectional Encoder Representations from Transformers (BERT) models using our datasets: (i) DNABERT pre-trained on the human whole genome^15^ and (ii) ViBE pre-trained on the viral genomes registered in the NCBI RefSeq Viral Database^16^ (**see Methods**). Given that the efficient replication and immune evasion of viruses are largely achieved through molecular mimicry of host organisms^3,17–19^, we hypothesized that the DNABERT model, pre-trained with the human genome, could extract representative features associated with human infectivity.

We evaluated model performance through benchmarking with existing models^3,6–10^. Candidate models were selected based on (i) their use of viral genome sequences as inputs and (ii) their applicability to several viral families (**Table 1).** In a preliminary experiment, the candidate models were re-trained on newly constructed datasets and compared with the original models (**Supplementary Fig. 1**). Notably, the three re-trained models, except for VIDHOP, outperformed their original counterparts on both the past and future virus datasets, as indicated by the improvement of both area under the receiver operating characteristic curve (AUROC) and area under the precision-recall curve (PR-AUC). These results highlight the contribution of our new datasets to enhancing the performance of viral infectivity prediction models.

**Table 1:** Predictive models for human infectivity using viral genetic information as inputs.

| Model name | Model architecture | Inputs or features | Model availability | In comparison |
| --- | --- | --- | --- | --- |
| BERT-infect<sub>DNABERT</sub> | DNABERT<sup>15</sup> | 250 bp subsequences | Fine-tuned DNABERT model<sup>42</sup> | Yes |
| BERT-infect<sub>ViBE</sub> | ViBE<sup>16</sup> | 250 bp subsequences | Fine-tuned ViBE model<sup>42</sup> | Yes |
| humanVirusFinder<sup>6</sup> | kNN | 4-mer frequency in >500 bp sequences | <sup>43</sup> | Yes |
| DeePac_vir<sup>7</sup> | CNN or LSTM | 250 bp subsequences | <sup>44</sup> | Yes |
| VIDHOP<sup>8</sup> | LSTM | 250 bp subsequences | <sup>45</sup> | No: due to the dataset size |
| HostNet<sup>9</sup> | Transformer-CNN-BiGRU | 250 bp subsequences | <sup>46</sup> | No: original models were unavailable |
| VirusBERTHP<sup>10</sup> | BERT | Sequences from 250 to 10,000 bp | <sup>47</sup> | No: original models were unavailable |
| zoonotic rank<sup>3</sup> | GBM | Viral genomic<br>feature and<br>similarity<br>feature to human<br>transcripts | 48 | Yes |

Abbreviations: kNN, k-nearest neighbor model; CNN, convolutional neural network model; LSTM, long short-term memory model; biGRU, bidirectional gated recurrent unit; GBM, gradient boosting machine.

### Performance comparison among models trained with the past virus datasets

To evaluate our model performance in predicting viral infectivity, we trained our and existing models using the past virus datasets with five-fold stratified cross-validation (**Figs. 2A-B**). BLASTn^20^ was used to assess the potential predictability of our datasets based on the simple hypothesis that the queried virus has the same infectivity with the most similar viral sequence (**see Methods**).

**Figure 2:**
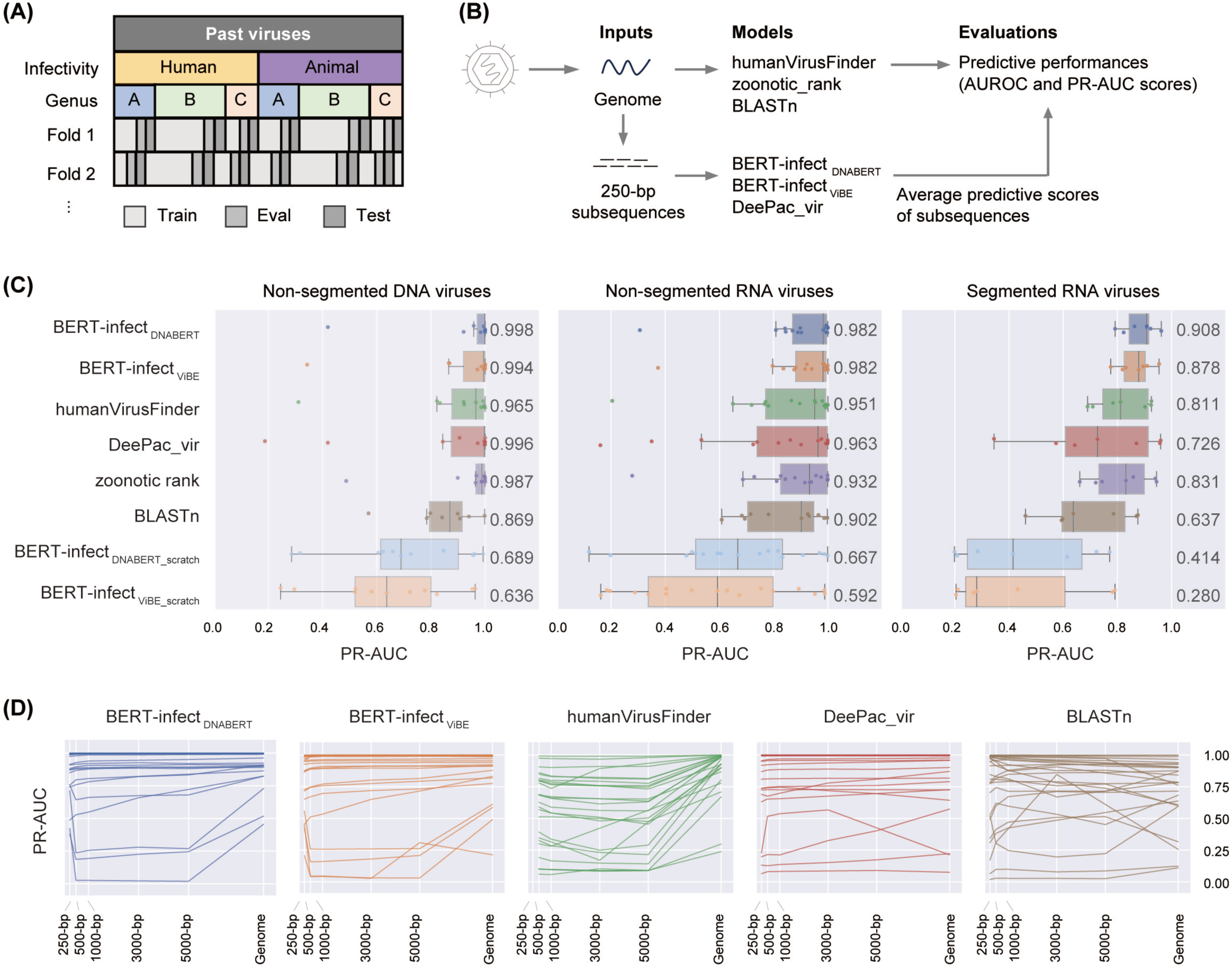
Predictive performance when using the past viral genomes as inputs. **a** Dataset division for five-fold stratified cross-validation. The fold datasets were prepared to maintain similar proportions of infectivity labels and viral genera as the overall data. **b** Differences in inputs and outputs of each model (**see Methods**). **c** Comparison of PR-AUC scores when inputting the past viral genomes. The median PR-AUC score is shown on the right side of the plot. Each dot corresponds to a viral family. **d** Changes in the PR-AUC score of each model according to the length of input sequences. Each line corresponds to a viral family. The 250 bp input is not available in the humanVirusFinder model, and its result is not included.

Our benchmark revealed that BERT-infect_DNABERT_ and BERT-infect_ViBE_ (fine-tuned with the past virus datasets) outperformed existing models across most viral families (**Fig. 2C**). Conversely, BERT-infect_DNABERT_scratch_ and BERT-infect_ViBE_scratch_ models (fine-tuned BERT models without pre-trained weights) failed to predict viral infectivity even with fine-tuning on the same datasets. The difference in performance between BERT-infect and previous models was especially pronounced when training with the relatively small datasets from the Virus-Host Database (**Supplementary Fig. 2A**). These results underscore the critical role of LLM pre-training for enhancing model performance, especially when labeled data is limited.

Remarkably, BERT-infect models demonstrated superior PR-AUC scores across 18 viral families, with particularly pronounced performance in the segmented RNA viruses. Despite comprising key pathogens associated with severe diseases, such as hemorrhagic fever^21^, these viral families have been neglected in previous model evaluations owing to data limitations (**Fig. 2C**). Thus, our comprehensive datasets represent a valuable resource for developing predictive models for viral infectivity across a range of viral families.

### Model applicability for zoonotic virus detection from high-throughput sequencing data

We evaluated model performance in detecting human-infecting viruses from high-throughput sequencing data when inputting (i) 250 bp single-ended high-throughput sequencing reads and (ii) viral contigs with various lengths, which reflect the real-world scenario where short viral contigs are often obtained using sequence assembly^22^. The BERT-infect and DeePac_vir models maintained consistent performance for most viral families, regardless of input length (**Fig. 2D**). We attributed the decrease in the predictive performance of the humanVirusFinder model to the k-mer compositions in short input sequence, which may not fully represent viral genome complexity^6^. These results indicated that only certain models are suitable for mining human-infecting viruses from high-throughput sequencing data.

We considered the computational resource required for model training and prediction (**Supplementary Table 7**). Deep learning-based models, including BERT-infect, can effectively process shorter input sequences, but they require considerable computational power and time. For example, fine-tuning the BERT-infect_DNABERT_ model with the Coronaviridae dataset takes 12 hours with four NVIDIA Tesla V100. In contrast, the humanVirusFinder and zoonotic rank models offer greater resource efficiency but are challenging to apply to high-throughput sequencing data. These observations underline the trade-offs between computational efficiency and model applicability to various types of inputs.

### Generalization performance in predicting infectivity of newly identified viruses

To provide early warnings on future pandemics, models should be able to predict the infectivity of newly discovered viruses. We evaluated the generalization capability of models trained on the past virus datasets when applied to the future virus datasets (**Fig. 1B**). For the family Coronaviridae, severe acute respiratory syndrome coronavirus 2 (SARS-CoV2)-related viruses (i.e., sarbecoviruses identified post-2018) were distinguished from other coronaviruses in the evaluation of model performance (**see Methods**).

Our approach considered two distinct thresholds (**Fig. 3A**). The first threshold was set based on the highest F1 scores from the past virus dataset analysis and served as a definitive criterion for distinguishing human-infecting viruses. We observed comparable median F1 scores in four models (**Fig. 3B**). In contrast, the DeePac_vir model displayed lower F1 scores, primarily due to decreased precision scores. However, it should be noted that the human infectivity of viruses in our dataset has not been experimentally validated, suggesting that some viruses may be mislabeled. Therefore, it is difficult to determine whether the low precision score is due to the high number of false positives or the detection of viruses unproven to be infectious to humans (**see Discussion, Supplementary Figs. 3-4**). For the second threshold, we explored the enrichment of human-infecting viruses among those predicted with the higher probabilities, which attempted to reflect a practical scenario where we need to prioritize high-risk viruses based on their predicted scores. We observed no notable differences across models; for instance, targeting viruses with the top 20% of predicted probabilities led to a median detection rate of ∼40% for human-infecting viruses (**Fig. 3C**). These results demonstrate a high level of model generalization toward novel viruses.

**Figure 3:**
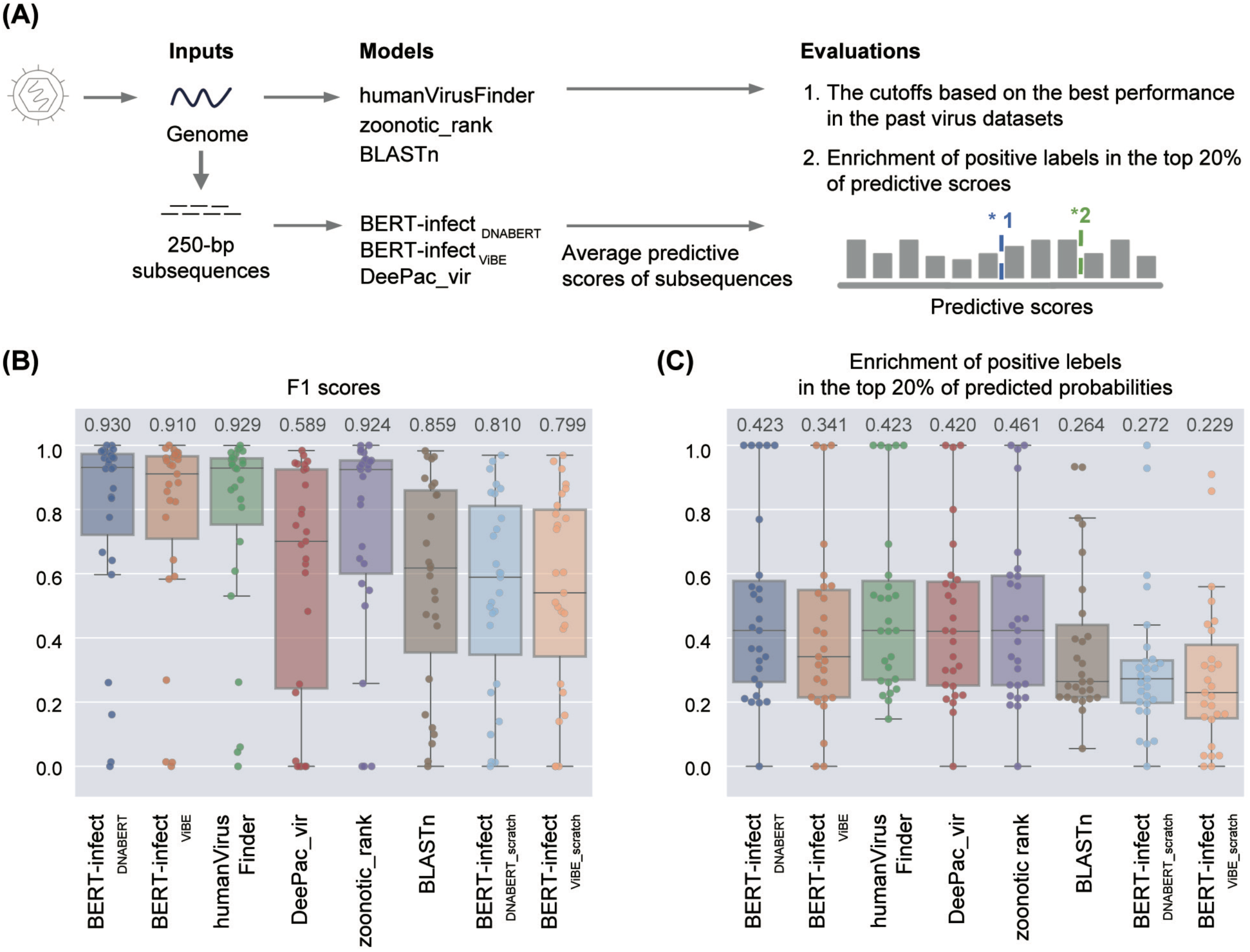
Generalization capabilities for predicting infectivity of future viruses. **a** Evaluation of model predictive performance for the future viral infectivity using two thresholds. **b** Model comparison when using the criteria representing the highest F1 scores in the past virus dataset. **c** The enrichment of human-infecting viruses in the top 20% of predicted probabilities. The y-axis shows the percentage of human-infecting viruses detected for each viral family in the top 20% of probabilities. The median score is shown above the plot.

### Systematic identification of difficult-to-predict viral lineages

To identify virus lineages for which human-infection risk is difficult to predict, we conducted a high-resolution model comparison at the viral family and genus levels (**Supplementary** Figs. 5-6). While the F1 scores were >0.75 for most virus families, poor predictive abilities were observed in almost all models for some viral families: Circoviridae, Coronaviridae, Flaviviridae, Hepeviridae, Rhabdoviridae, and Hantaviridae. Furthermore, even when F1 scores were above 0.75 for other viral genera, there were predictive gaps within particular viral genera: SARS-CoV2-related viruses, Flavivirus, and Phlebovirus, associated with severe infectious diseases and suspected zoonotic origins. Such difficulties were also observed in the original model, although not noted in previous studies^3,6,7^. Notably, our analysis revealed that no current models could adequately warn of the human infection risk posed by SARS-CoV2-related viruses (**details in the subsequent section**). Our findings highlight critical issues in viral infectivity prediction models: despite achieving substantial predictive performance overall, the models consistently failed to predict the infectivity of specific zoonotic viral lineages.

### Difficult-to-predict viral lineage frequently changed infectivity during evolution

To further investigate the challenges in predicting human infectivity of zoonotic viruses, we mapped the phylogenetic relationships of viruses alongside their prediction results (**Fig. 4A**). Here, we focused on Flaviviridae, which includes viruses that cause severe infectious diseases in humans, such as hepatitis C virus (genus Hepacivirus) and Zika virus (genus Flavivirus). In the genus Hepacivirus, a phylogenetic distinction existed between human- and animal-infecting viruses, and F1 scores for all models were above 0.8 for both the past and future virus datasets (**Fig. 4B**). In contrast, model performance dropped for the genus Flavivirus, which was characterized by frequent shifts in infectivity during its evolution: the F1 scores were ∼0.6 for the future virus dataset. These results emphasize the insufficient generalization capability of current models for viral lineages where the zoonotic-infection potential has been acquired repeatedly.

**Figure 4:**
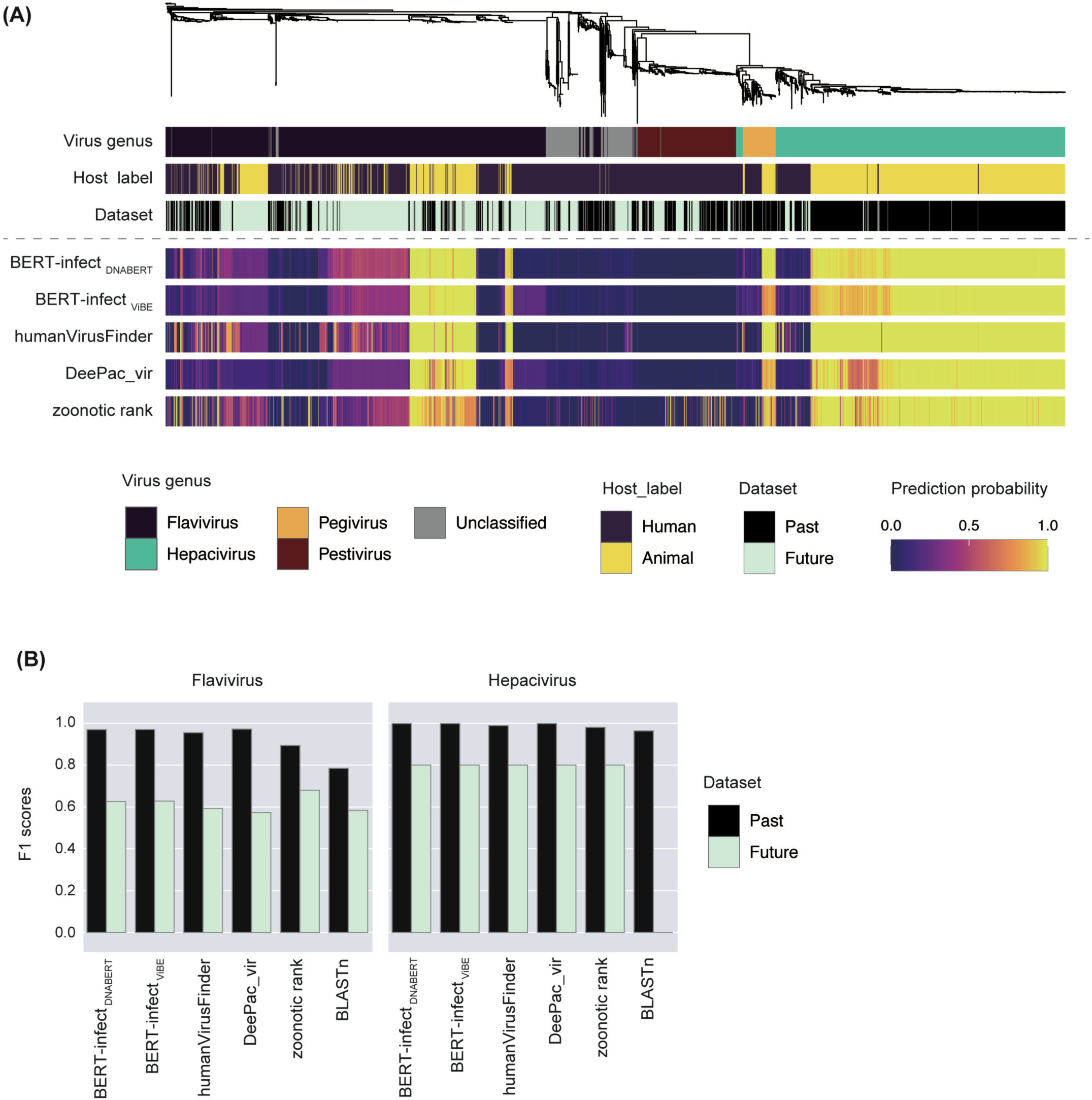
Predictive viral infectivity for Flaviviridae. **a** Association between viral phylogenetic relationships and predicted results. The upper panel shows the phylogenetic tree, and the heatmaps in the middle panel show each viral property (i.e., viral genus, host label, and dataset). The lower panel shows the predictive probabilities generated by the five models. **b** Comparison of the F1 scores among viral genera.

### Challenges in predicting SARS-CoV-2-related viral infectivity

Our study revealed severe limitations in the ability of models to identify the risk of human infectivity posed by SARS-CoV2-related viruses (**Supplementary Fig. 5**). All models trained on the past virus dataset failed to recognize SARS-CoV2-related viruses as a potential threat despite achieving high predictive performance for most coronaviruses (**Fig. 5A-B**). Notably, although this dataset included various human-infecting SARS-CoV-2 variants from the original outbreak and the currently prevalent Omicron variants, the human-infecting potential of SARS-CoV2-related viruses could not be determined based on the best F1 score for the past virus datasets (**Fig. 5C**). These findings highlight a crucial gap in our preparedness for zoonotic pandemics: even if SARS-CoV2-related viruses had been detected before the COVID-19 outbreak through genomic surveillance, the current models would not have flagged them as high-risk (**see Discussion**).

**Figure 5:**
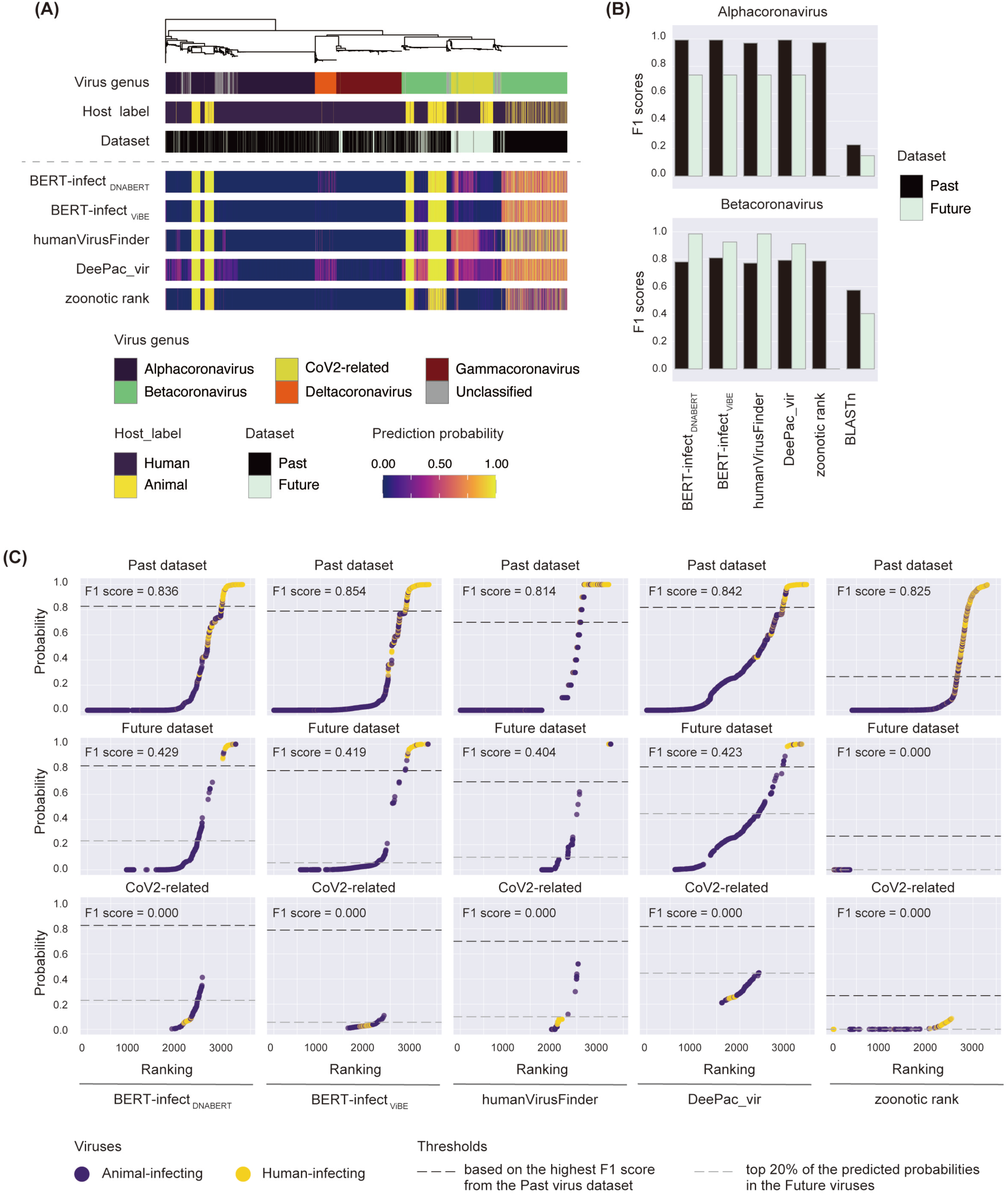
Predictive viral infectivity for Coronaviridae. **a** Association between viral phylogenetic relationships and predicted results. The upper panel shows the phylogenetic tree, and the heatmaps in the middle panel show each viral property (i.e., viral genus, host label, and dataset). The lower panel shows the predictive probabilities generated by five models. **b** Comparison of the F1 scores among viral genera. **c** Ranking of predictive probabilities in the past virus dataset (upper), future virus dataset (middle), and SARS-CoV2-related viruses (bottom).

## Discussion

The rapid elucidation of viral genetic diversity has increased the demand for high-throughput approaches for assessing the potential human infectivity of viruses, such as machine learning models using viral genetic information as inputs. In this study, to overcome the limitations of insufficient labeled data in constructing viral infectivity prediction models, we constructed new datasets across 26 viral families and developed innovative models utilizing pre-trained LLMs for context-like rules within genetic sequences (**Fig. 1**). Our models exhibited high predictive performance for the past and future virus datasets across most viral families (**Figs. 2-3**). Particularly noteworthy was the improved performance of our models on segmented RNA viruses, which have been neglected in viral infectivity prediction owing to limited data. These results represent a major advance in our preparedness to combat the future threats of zoonotic viruses.

While models trained on the past virus datasets demonstrated high generalization capabilities for most viruses that emerged after model construction, our high-resolution evaluation based on phylogenetic analysis revealed significant limitations in current models for assessing the risk associated with specific zoonotic viral lineages (**Figs. 4-5, Supplementary Fig. 5**). Such limitations were observed even in models previously reported to have high predictive performance, indicating an essential area for future development that has not been addressed before. Crucially, no model has accurately identified the human infection risk posed by SARS-CoV2-related viruses, and previous studies have also only weakly warned of the risks associated with these viruses. Therefore, even with enhanced viral surveillance in animal populations, current models may fail to detect emerging high-risk zoonotic viruses. Our insights emphasize the need for advancements in predictive models to prepare against future zoonotic viral diseases.

A potential limitation in developing models to predict viral infectivity is the gap between the predictive purpose and the training datasets, where the labels may not accurately reflect the actual viral infectivity. Typically, most models are trained using infectivity labels defined based on host information from which organism the viral sequences were identified (**Fig. 1A**). This is an alternative approach due to the limited availability of comprehensive datasets on viral infectivity in humans^6–10^. Thus, it should be noted that a certain number of human-infecting viruses may be labeled as animal ones. Given such limitations, animal-derived viruses predicted to be human-infectious by multiple models should be prioritized for risk assessment in future research (**Supplementary Fig. 4**).

In this study, we constructed new datasets across 26 viral families, comprising ∼29-times more data available than previously used^14^ (**Fig. 1C**); however, further dataset curation is necessary to enhance the model performance of zoonotic viral infectivity prediction. The exclusive inclusion of easy-to-predict viruses could hinder learning for zoonotic viruses derived from rare spillover events^2,12,13^. Our evaluation showed high generalization capabilities for most viral families in the future virus datasets, likely owing to the presence of viruses having high sequence similarity and the same infectivity as the past viruses (**Supplementary Fig. 3**). Thus, further dataset curation focusing on viral lineages associated with zoonoses could enhance generalization performance. However, it should also be noted that removing redundancy involves a trade-off with the scarcity of data, because a limited number of human infectious viruses have been experimentally validated.

Furthermore, another limitation in our dataset curation is that the data labeling method cannot distinguish between infectivity (i.e., capability for animal-human infection) and transmissibility (i.e., ability to expand human-human infections). In this study, we also conducted large-scale predictions on the infectivity of H5Nx influenza A viruses, which caused zoonotic spillovers after 2018 (**Supplementary Fig. 6**). Still, current models did not predict most zoonotic spillover viruses as high risk. On the other hand, it is currently difficult to determine whether these results reflect that recent spillover events by the H5Nx influenza A viruses have not led to human-to-human transmission or due to the inadequate predictive performance of the current models. Further model developments to hierarchically predict infectivity and transmissibility would be necessary for accurately assessing the risk of zoonotic spillover and subsequent pandemic potential. Models using nucleotide sequences as inputs may effectively detect host adaptations in viral genome compositions; however, capturing infectivity changes due to minor genomic variations remains challenging. Indeed, our evaluation revealed that current models fail to recognize the human infectivity threat posed by SARS-CoV2-related viruses (**Fig. 5**). One explanation is infectivity changes driven by limited mutations, mainly in the spike protein^23,24^. Utilizing protein language models, known for extracting feature vectors that reflect the structural and functional properties of proteins^11^, could enhance model performance by detecting key changes at the amino acid level relevant to viral infectivity^25^.

Knowledge-based models integrating multiple features related to molecular mechanisms underlying viral infectivity represent another promising approach^26^. Recent studies highlighted the importance of various types of virus-host interactions, beyond the entry pathway, in determining viral infectivity^3,17,27^. Interestingly, our BERT-infect models displayed comparable predictive performance, although the DNABERT and ViBE models were pre-trained on the human and viral genomes, respectively (**Figs. 2-3, Supplementary Fig. 5**). These results suggest that two BERT-infect models may capture different contextual features involved in viral infectivity and yield new insights into viral infection mechanisms. Thus, a dual strategy of (i) model refinement for unsupervised feature extraction and (ii) deepened understanding of infection mechanisms through model interpretation would contribute to developing knowledge-based models for accurately assessing the human infection risk of viruses.

In conclusion, this study presents new models leveraging LLMs along with a framework for evaluating model performance in predicting viral infectivity. Although our models demonstrated high predictive performance for various viral families, we found unresolved challenges in accurately predicting certain zoonotic viral lineages across the current models. Our findings provide valuable perspectives on enhancing the generalization capabilities of predictive models, contribute to developing a risk assessment tool against potential zoonotic spillover threats.

## Methods

### Preparing viral sequences and infectivity datasets

Viral sequences and metadata for the 26 viral families were collected from the NCBI Virus Database^28^ (**Fig. 1A**). Viral infectivity was labeled according to the host information, from which organism the viral sequences were isolated, and the human infectivity was not experimentally verified for all viruses. For segmented RNA viruses, which include multiple sequences, we grouped sequences into viral isolates based on the combination of 22 entries in the metadata (**Supplementary Table 2**). If a single viral isolate contained more sequences than the specified number of viral segments (i.e., Orthomyxoviridae: 8, Rotaviridae: 12, and other segmented RNA viruses: 3), redundancy was eliminated by randomly sampling two sequences for each segment. For non-segmented viruses, representative sequences were selected from identical viral sequences with the same infectivity label. Our curation strategy generated ∼29-times more viral data available than that of the Virus-Host Database used predominantly in previous studies (**Supplementary Table 1**).

We divided the collected data into a past virus dataset for model training and a future virus dataset for evaluating model generalization (**Fig. 1B**). Data with a sequence collection date before December 31, 2017, were classified into the past virus datasets, while the subsequent data were classified into the future virus datasets. To validate the applicability of viruses associated with repeated zoonoses, we collected additional future virus dataset for Orthomyoviridae and Coronaviridae families from several different resources: (i) Influenza A viruses classified into four H subtypes (i.e., H1, H3, H5, and H7) from the NCBI Influenza virus database^29^, (ii) Influenza A virus classified into H5 subtype from the GISAID database(https://gisaid.org/), and (iii) SARS-CoV2-related viruses (i.e., sarbecoviruses identified after 2018) with animal infectivity from previous study^30^ and those with human infectivity from the Nextstrain database^31^ (**Supplementary Tables 3-4**). For the GISAID dataset prediction, we annotated the position of open reading frame (ORF) by EMBOSS getorf (version 6.6.0.0)^32^ and used them as inputs of the zoonotic rank model.

### Constructing predictive model for viral infectivity based on LLMs

We constructed new predictive models for viral infectivity, namely BERT-infect, based on the LLMs. We used the (i) DNABERT model pre-trained on human whole genome^15^ and (ii) ViBE model pre-trained on the viral genome sequences in the NCBI RefSeq database^16^. Since pre-trained models with 4-mer tokenization were available in both the DNABERT and ViBE models, we fine-tuned these BERT models using the past virus datasets to construct an infectivity prediction model for each viral family. Input data were prepared by splitting viral genomes into 250 bp fragments with a 125 bp window size and 4-mer tokenization. The hyperparameters used to fine-tune two BERT models are listed in **Supplementary Table 5**.

### Re-training comparative models

Four existing models (humanVirusFinder^6^, DeePac_vir^7^, VIDHOP^8^, and zoonotic_rank^3^) were re-trained with our newly constructed datasets (**Supplementary Fig. 1**). The hyperparameters for the DeePac_vir and VIDHOP model re-training are listed in **Supplementary Table 6**. The humanVirusFinder and zoonotic_rank models were re-trained with the same parameters as in the previous studies. For humanVirusFinder model, we used the 4-mer frequency as inputs, which exhibited best performance in previous study. The zoonotic rank model was re-trained using all genome composition features (i.e., viral genomic features, similarity to interferon-stimulated genes, similarity to housekeeping genes, and similarity to remaining genes) over 1,000 iterations, and the top 10% of iterations were used to construct the bagged model.

In a preliminary experiment, we compared predictive performance between the re-trained and original models (**Supplementary Fig. 1**). Three re-trained models outperformed their original counterparts on the past and future virus datasets. In contrast, the original VIDHOP model showed higher predictive performance than the re-trained model, likely due to its original training on >10,000 sequences per viral family^8^. Besides, the original HostNet^9^ and VirusBERTHP^10^ models, trained on similar-sized datasets as those of VIDHOP, were unavailable and excluded from the subsequent analyses.

We used BLASTn^20^ to evaluate potential dataset predictability based on the simple hypothesis that a virus has the same infectivity as that with the highest sequence similarity. BLASTn prediction was conducted using the training dataset as a database and test dataset as a query. The following cases were judged as unpredictable: (i) there was no hit sequence with an E-value of >1e-4, (ii) a viral sequence showed both human- and animal-infecting viruses with the same bitscore, or (iii) the aligned length of the top hit sequence did not cover >50% of the query sequence.

### Evaluation of viral infectivity prediction for the past viral genome

In model training and validating using the past virus datasets, stratified five-fold cross-validation was performed to adjust for the class imbalance of infectivity and virus genus classifications (**Fig. 2A**). The training, evaluation, and test datasets proportions were set to 60%, 20%, and 20%, respectively. The prediction probabilities were calculated in differently for each type of model. The humanVirusFinder and zoonotic_rank models use the viral genome sequence as inputs and directly output the predicted results for each sequence. In contrast, the BERT-infect, DeePac_vir, and VIDHOP models use 250 bp subsequences as inputs, and the prediction results for genomic sequences were calculated by averaging the predicted scores for subsequences (**Fig. 2B**). The predictive performances were compared using two metrics: (i) the area under the receiver operating characteristic curve (AUROC) and (ii) the area under the precision-recall curve (PR-AUC). Down-sampling was conducted for virus families with PR-AUC <0.75 in all models when training with the original past virus dataset (i.e., Rhabdoviridae, Circoviridae, Poxviridae, and Hantaviridae).

### Evaluation of model detection ability for human-infecting virus using high-throughput sequencing data

We evaluated model predictive performance when using high-throughput sequencing data based on different inputs: (i) 250 bp single-ended high-throughput sequence reads and (ii) variable-length contig sequences (i.e., 500 bp, 1,000 bp, 3,000bp, and 5,000 bp) (**Fig. 2D**). For the first input, we only compared BERT-infect_DNABERT_, BERT-infect_ViBE_, and DeePac_vir because the humanVirusFinder and zoonotic_rank models have a recommended input of >500 bp sequences and viral genomic sequences, respectively. For the second input, we compared BERT-infect_DNABERT_, BERT-infect_ViBE_, humanVirusFinder, and DeePac_vir. The humanVirusFinder model directly outputted the predicted results for each contig, whereas the prediction results of the other three models were calculated by averaging the predicted scores for subsequences (**Fig. 2B**). We evaluated model performance based on the AUROC and PR-AUC.

### Evaluation of viral infectivity prediction for the future viral genome

Model predictive performance for the future viral dataset was evaluated under the two scenarios (**Fig. 3A**). First, the infectivity of novel viruses was determined based on the threshold representing the highest F1 score in the past viral datasets, and F1, recall and precision metrics were calculated (**Fig. 3B**). Second, we investigated the enrichment of human-infecting viruses within the top 20% of predicted probabilities (**Fig. 3C**).

### Phylogenetic analyses

Multiple sequence alignments for each viral family were constructed by adding sequences to the reference alignment provided by the International Committee on Taxonomy of Viruses resources^33^. First, we eliminated sequence redundancy in our datasets using CD-HIT (version: 4.8.1)^34,35^ when the number of sequences per label was >200. For nucleotide reference sequences, our dataset sequences were added into the reference alignment using mafft (version: 7.508)^36^ with the options “--add” and “--keeplength”. For amino acid reference sequences, the protein sequences were extracted from viral genomic sequences using tBLASTn (version: 2.15.0)^20^ and were added into the reference alignment using mafft with the options “--add” and “--keeplength”. Phylogenetic trees were constructed by the maximum likelihood method using IQ-TREE (version 2.1.4 beta)^37^. Substitution models were selected based on the Bayesian information criterion provided by ModelFinder^38^. The branch supportive values were measured using ultrafast bootstrap in UFBoot2^39^ with 1,000 replicates. Visualization of the phylogenetic tree, viral characteristics, and their prediction results were performed using ggtree package (version 3.6.0)^40^. The parameters used to construct the phylogenetic tree for each viral family are listed in **Supplementary Table 8**.

### Data availability

The data used in this study were published in the Zenodo Repository^41^.

### Code availability

The relevant codes are available at https://github.com/Junna-Kawasaki/BERT-infect_2023.

## Acknowledgment

We acknowledge with gratitude the Authors, the Originating laboratories responsible for obtaining specimens, and the Submitting laboratories that generated and shared the genetic sequence and metadata via the GISAID Initiative. We are grateful to Dr. Jumpei Ito (The University of Tokyo, Japan), Dr. Sho Miyamoto (National Institute of Infectious Diseases, Japan), and Dr. Masayuki Horie (Osaka Metropolitan University, Japan) for their helpful discussions. This study was supported by grant from JST PRESTO (JPMJPR23R4 to J.K.); Japan Society for the Promotion of Science (JSPS) KAKENHI (JP22KJ2901 to J.K.); by the Waseda Research Institute for Science and Engineering, Grant-in-Aid for Young Scientists (Early Bird) (J. K.). The super-computing resource was provided by the ROIS National Institute of Genetics, the Human Genome Center at The University of Tokyo, and the Information Technology Center at The University of Tokyo. We thank Editage (https://www.editage.com/) for editing and reviewing this manuscript for English language. Some illustrations were retrieved from BioRender.com.

## Author contributions

J.K. and M.H. conceived the study; J.K. mainly performed the bioinformatics analyses;

J.K. prepared the figures and wrote the initial draft of the manuscript. All authors contributed to study design, data interpretation, and paper revision, and have approved the final manuscript for publication.

## Materials & Correspondence

Correspondence and material requests should be addressed to Junna Kawasaki or Michiaki Hamada

## Supplementary information

**Supplementary Figure 1:**
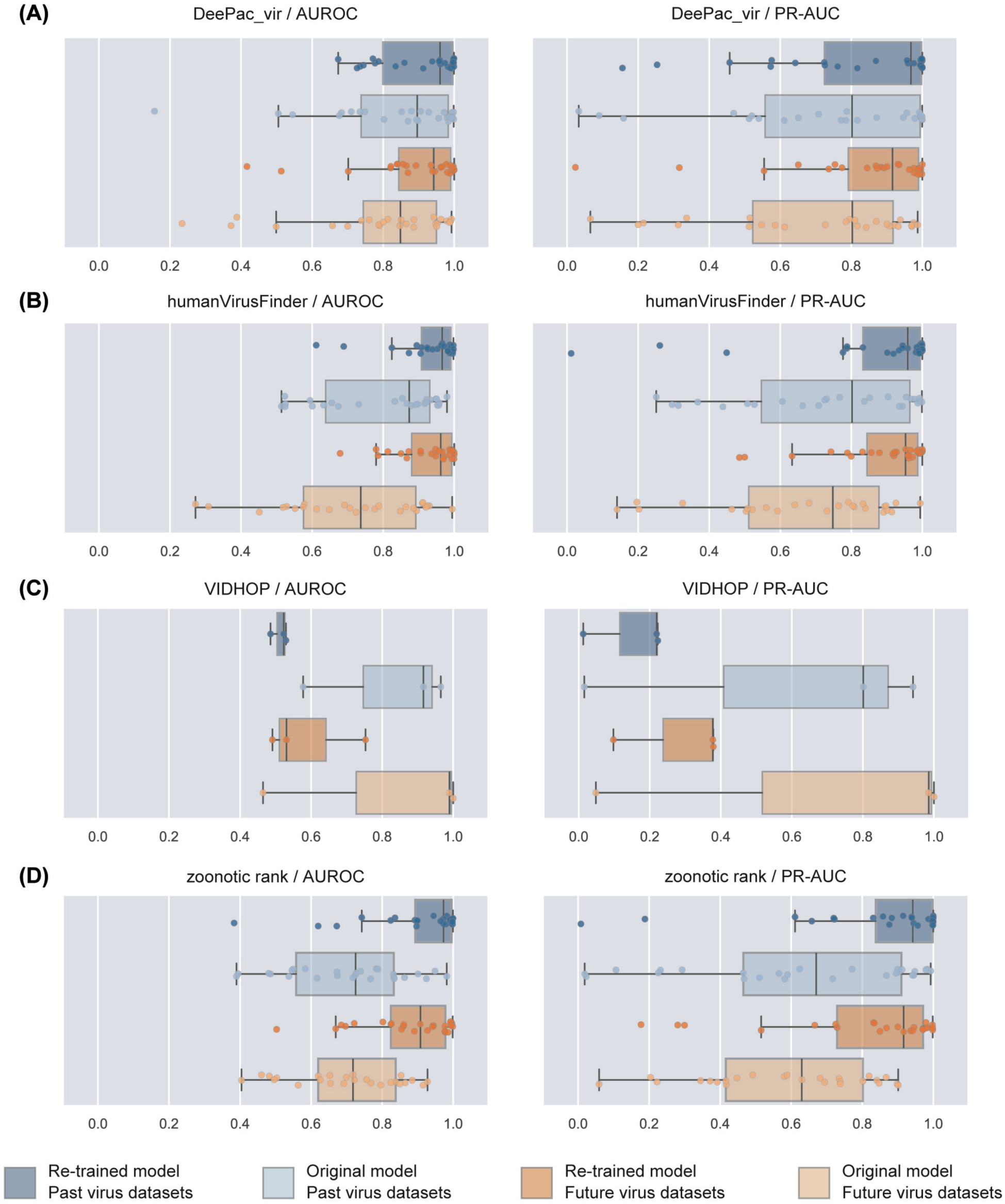
Comparison between re-trained and original models. **a–d** AUROC (left) and PR-AUC (right) scores of the re-trained and original models: **a** DeePac_vir, **b** humanVirusFinder, **c** VIDHOP, and **d** zoonotic rank. The graph color corresponds to the combination of the model and prediction dataset. The original VIDHOP model was constructed for only three viral families: Orthomyxoviridae, Reoviridae, and Rhabdoviridae. It should be noted that data leakages for training datasets of the original models were not prevented.

**Supplementary Figure 2:**
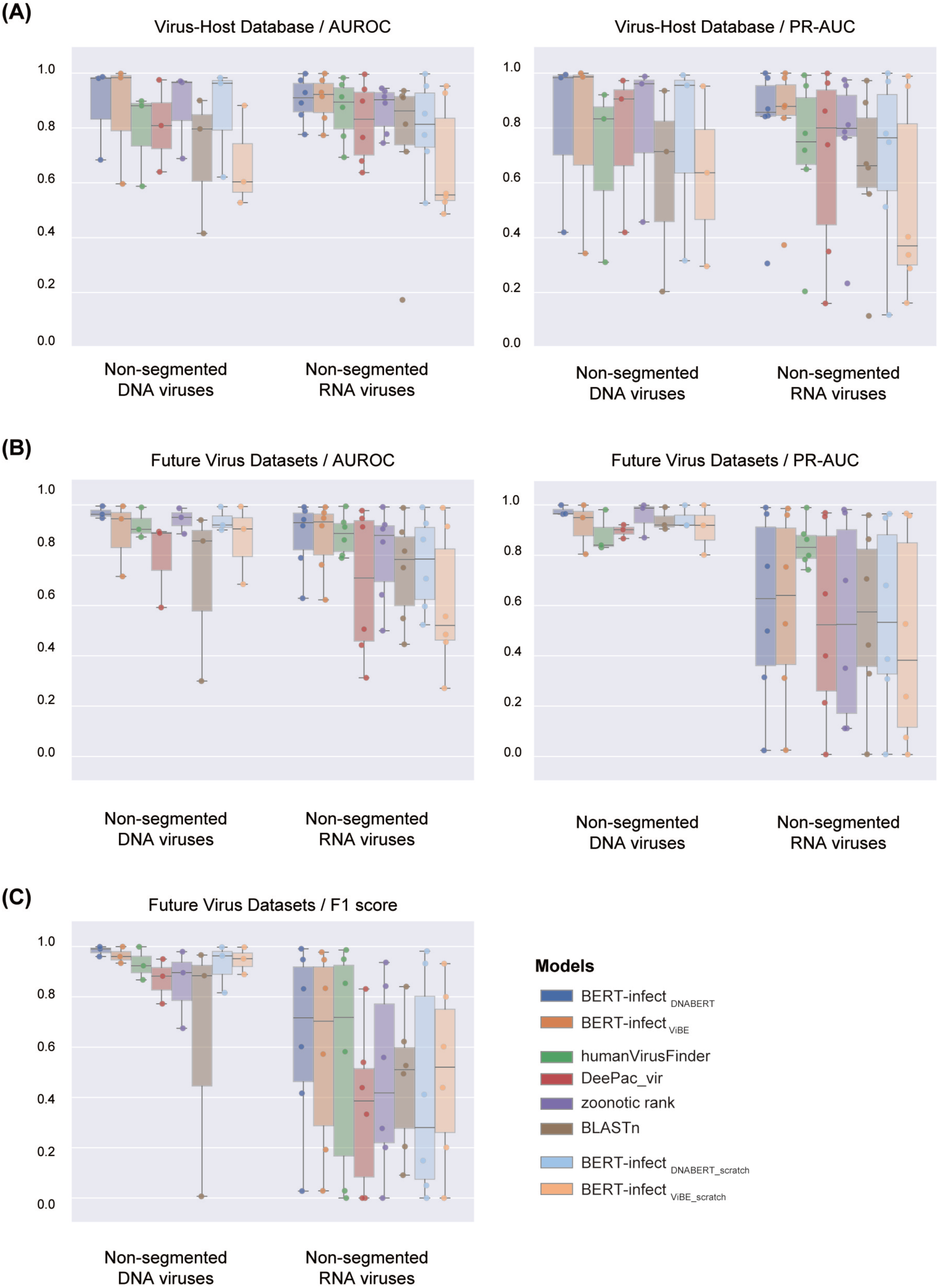
Predictive performance of models trained on the Virus-Host Database. Comparison of the AUROC (left) and PR-AUC (right) scores when **a** using the Virus-Host Database for model training and **b** using the future viral datasets for prediction. **c** F1 scores in future virus datasets using the threshold representing the highest F1 scores in the Virus-Host Database. We evaluated three DNA viruses (Adenoviridae, Hepadnaviridae, and Poxviridae) and six RNA viruses (Caliciviridae, Coronaviridae, Flaviviridae, Paramyxoviridae, Picornaviridae, and Rhabdoviridae). Each dot corresponds to a viral family, and the x-axis is categorized based on the viral genomic type.

**Supplementary Figure 3:**
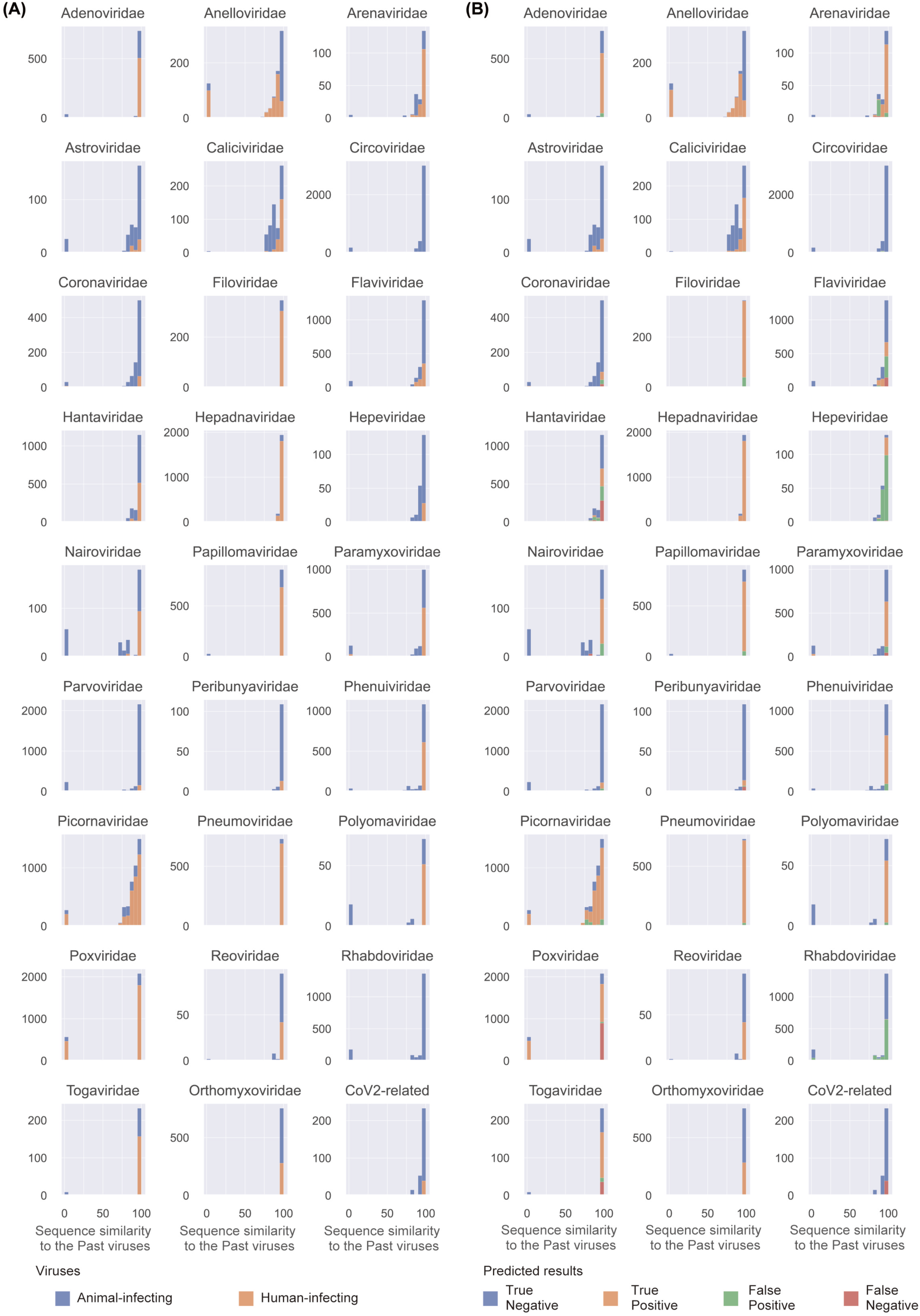
Association between predicted results for the future viruses and their sequence similarity with the training datasets. **a** Distribution of sequence similarity between the future viruses and training datasets for each viral family. **b** BERT-infect_DNABERT_ prediction results for future viruses per sequence similarity to the training datasets. The prediction results were determined based on the threshold representing the highest F1 scores in the past virus datasets.

**Supplementary Figure 4:**
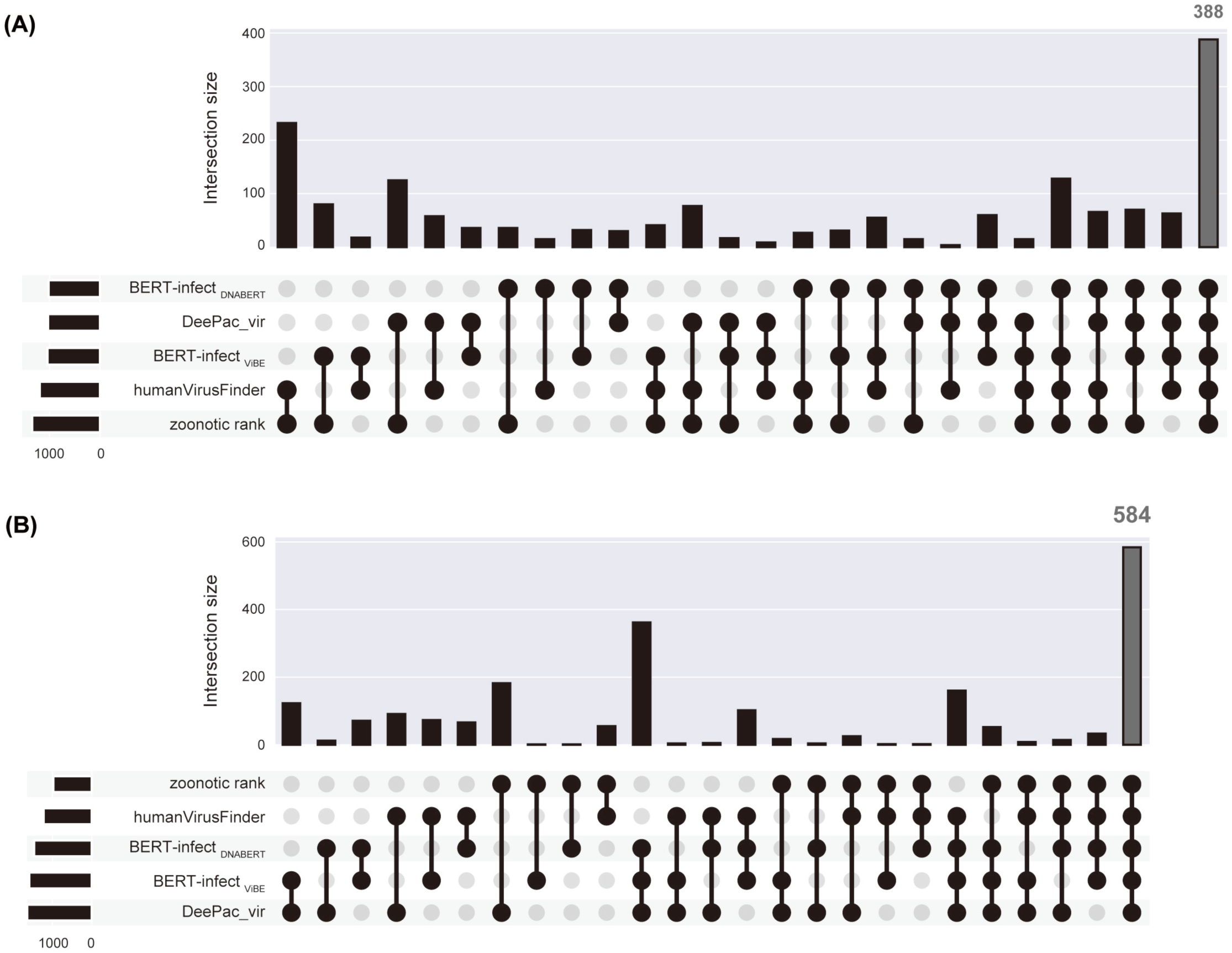
Animal-derived viruses with suspected human infectivity. Number of viruses predicted as potentially human infectivity in **a** past and **b** future viral datasets.

**Supplementary Figure 5:**
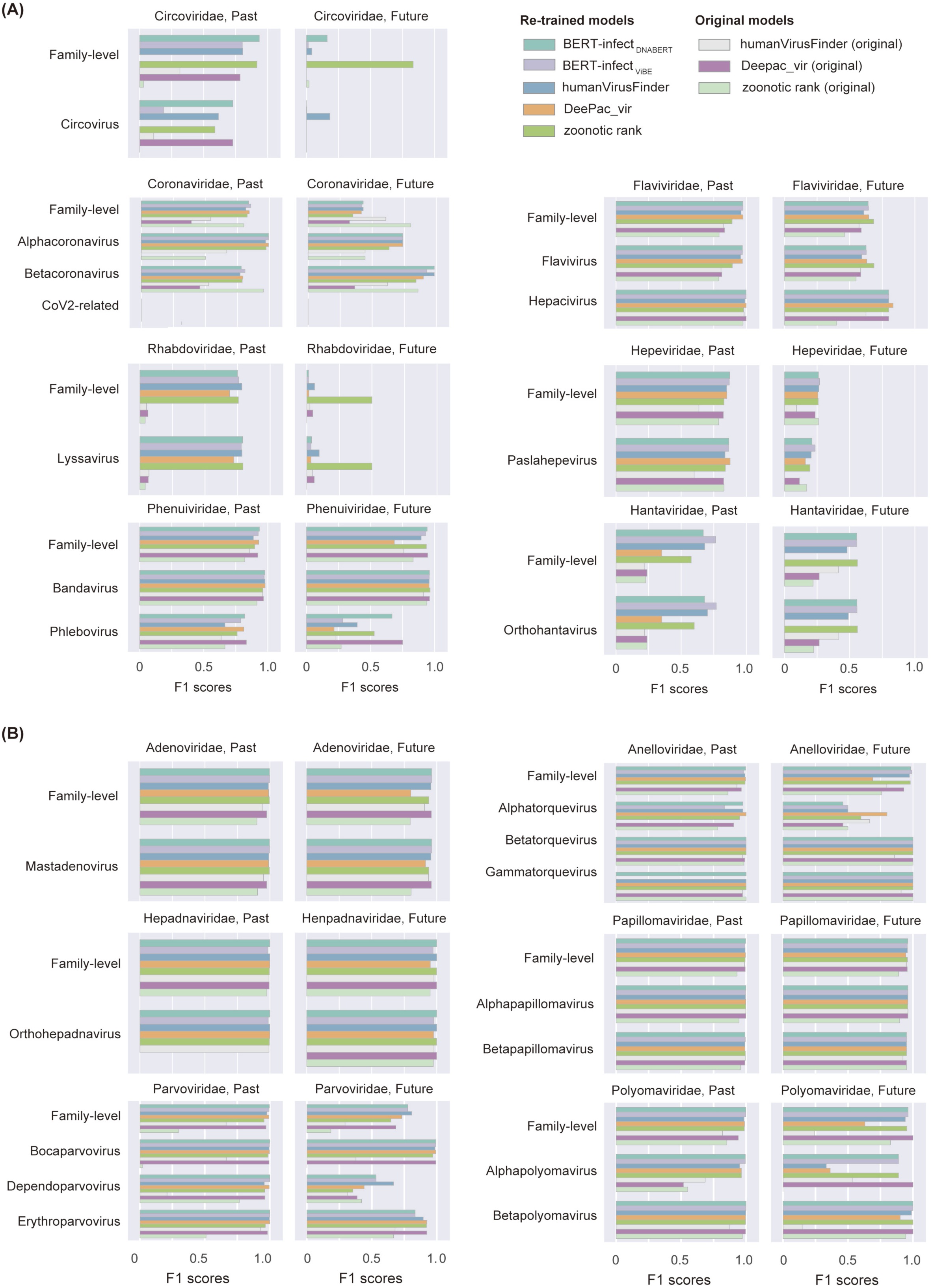
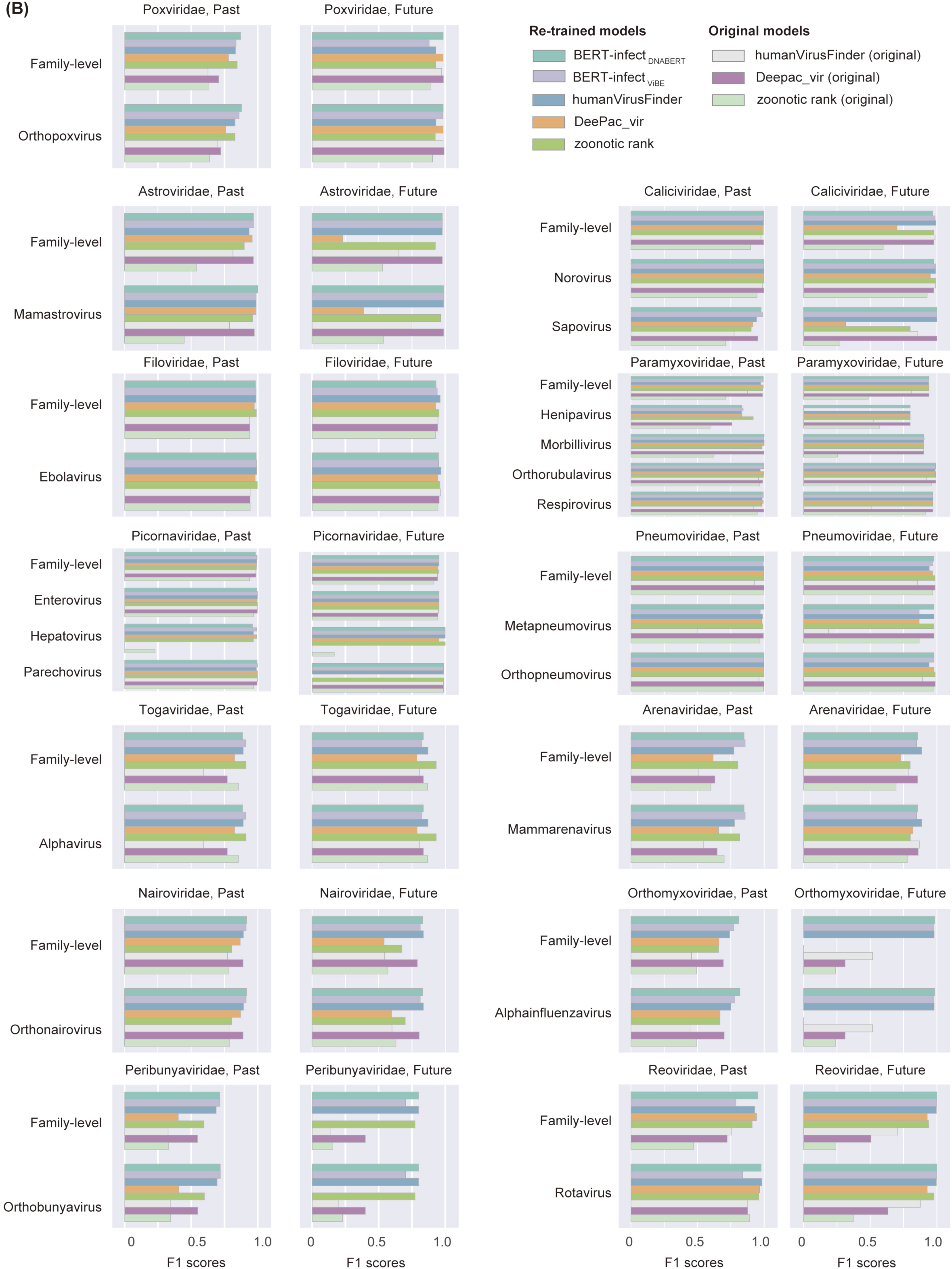
Comparison of F1 scores among viral families/genera. F1 scores below 0.75 at the viral family and genus level for almost all models are plotted in **a**, otherwise in **b**. Virus genera that met the following were plotted: (i) include viruses that infect both humans and animals, (ii) the past virus dataset contains >30 viral strains, of which >10% are human viruses, and (iii) the future virus dataset contains >10 viral strains, with >3 human-infectious viruses.

**Supplementary Figure 6:**
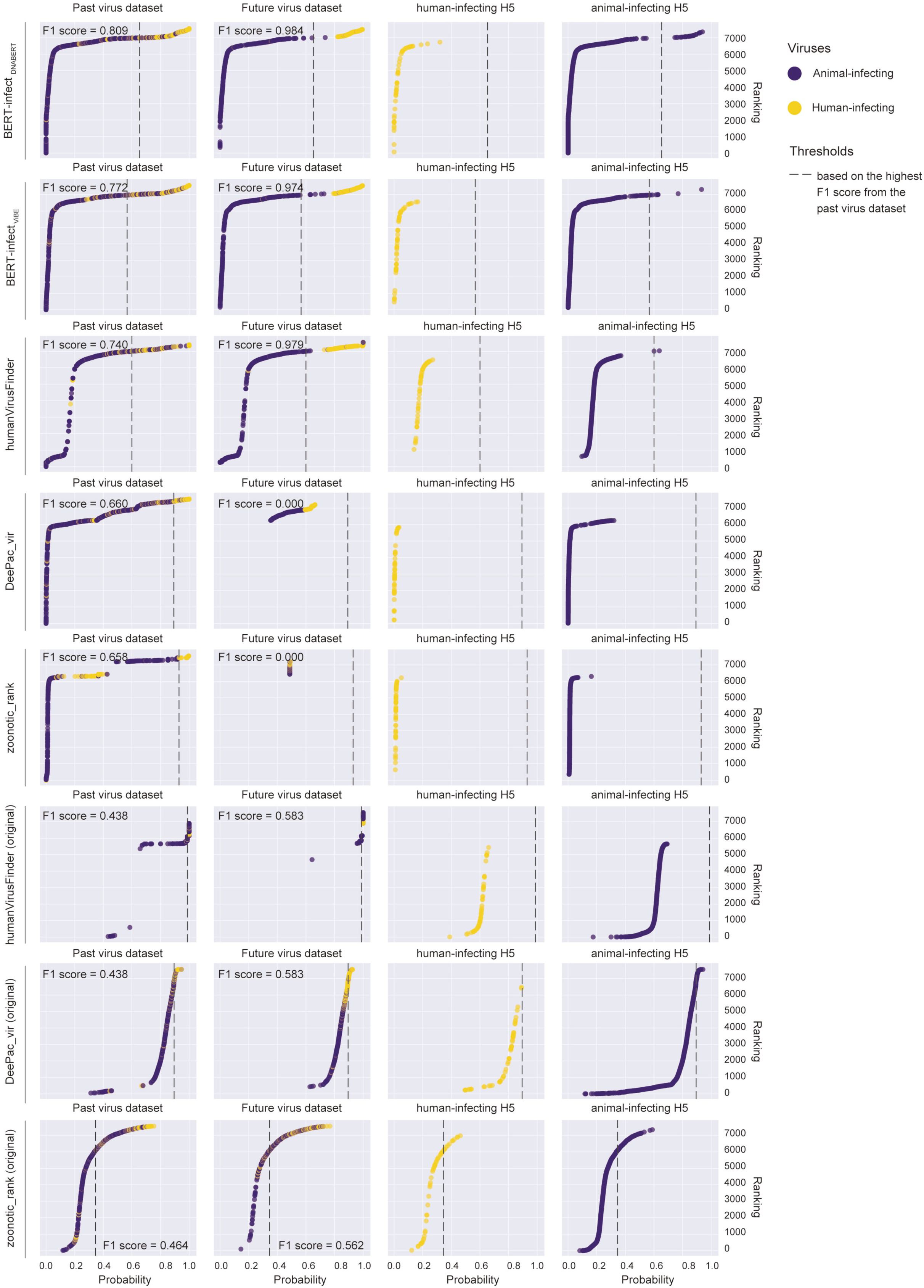
Prediction of zoonotic spillover risk for Influenza A virus subtype H5Nx. Ranking of predictive probabilities in the datasets of past viruses (left), future viruses (middle), and Influenza A virus subtype H5 (right).

**Supplementary Table 1: Dataset obtained from the Virus-Host Database**

**Supplementary Table 2: Our datasets for 26 viral families collected from the NCBI Virus Database**

**Supplementary Table 3: Datasets of Influenza A viruses identified after 2018**

**Supplementary Table 4: Datasets of SARS-CoV-2-related viruses identified after 2018**

**Supplementary Table 5: Hyperparameters used in fine-tuning of BERT models**

**Supplementary Table 6: Re-training conditions for the comparative models (DeePac_vir and VIDHOP)**

**Supplementary Table 7: Computational resources and time for model construction and infectivity prediction**

**Supplementary Table 8: Parameters for constructing the phylogenetic tree for each virus family**

